# Structural basis of V-ATPase V_O_ region assembly by Vma12p, 21p, and 22p

**DOI:** 10.1101/2022.10.19.512923

**Authors:** Hanlin Wang, Stephanie A. Bueler, John L. Rubinstein

## Abstract

Vacuolar-type ATPases (V-ATPases) are rotary proton pumps that acidify specific intracellular compartments in almost all eukaryotic cells. These multi-subunit enzymes consist of a soluble catalytic V_1_ region and a membrane-embedded proton-translocating V_O_ region. V_O_ is assembled in the endoplasmic reticulum (ER) membrane and V_1_ is assembled in the cytosol. However, V_1_ binds V_O_ only after V_O_ is transported to the Golgi membrane, thereby preventing acidification of the ER. We isolated V_O_ complexes and subcomplexes from *Saccharomyces cerevisiae* bound to V-ATPase assembly factors Vma12p, Vma21p, and Vma22p. Electron cryomicroscopy shows how the Vma12-22p complex recruits subunits a, e, and f to the rotor ring of V_O_ while blocking premature binding of V_1_. Vma21p, which contains an ER-retrieval motif, binds the V_O_:Vma12-22p complex, ‘mature’ V_O_, and a complex that appears to contain a ring of loosely-packed rotor subunits and the proteins YAR027W and YAR028W. The structures suggest that Vma21p binds assembly intermediates that contain a rotor ring, and that activation of proton pumping following assembly of V_1_ with V_O_ removes Vma21p, allowing V-ATPase to remain in the Golgi. Together, these structures show how Vma12-22p and Vma21p function in V-ATPase assembly and quality control, ensuring the enzyme acidifies only its intended cellular targets.

## Introduction

Vacuolar-type adenosine triphosphatases (V-ATPases) are membrane-embedded enzyme complexes that couple ATP hydrolysis to proton translocation across a membrane ^1^. V-ATPases are ubiquitous in eukaryotic cells where they acidify intracellular compartments such as vacuoles, lysosomes, endosomes, and the trans Golgi network ^2,3^. V-ATPases in the plasma membrane of specialized cells, including osteoclasts ^4^, kidney intercalated cells ^5^, and some cancer cells ^6^, acidify the extracellular environment. Due to their numerous roles, V-ATPase malfunction can lead to various diseases such as osteopetrosis ^7,8^, distal renal tubular acidosis ^9^, and neurodegenerative disorders ^10,11^.

V-ATPases consist of a soluble V_1_ region that hydrolyzes ATP and a membrane-embedded V_O_ region that translocates protons ^1,3^. Within V_1_, a hexamer of subunits A and B catalyzes ATP hydrolysis, causing conformational changes that drive rotation of a central rotor (subunits DFd) that is attached to a membrane-embedded c ring (subunits c_8_, c’, c”, and Voa1p) within the V_O_ region ^12–14^. Rotation of the c ring against the membrane-embedded C-terminal domain of subunit a drives proton translocation. The membrane-embedded subunits e and f, which have unknown functions, bind the C-terminal domain of subunit a. During rotation of the c ring, protons enter a cytosolic half channel between subunit a and the c ring, protonating conserved glutamate residues on each of the c, c’, and c” subunits. Continued rotation of the ring carries protons through the lipid bilayer to a luminal half channel in subunit a where they exit the complex on the other side of the membrane ^12^. Three pairs of E and G subunits form peripheral stalk structures that interact with the soluble subunits C and H, as well as the cytosolic N-terminal domain of subunit a, holding subunit a stationary relative to the rotating c ring. As a result of its rotary catalytic mechanism, electron cryomicroscopy (cryoEM) of intact V-ATPase shows the enzyme in three main conformations, known as rotational ‘State 1’, ‘State 2’, and ‘State 3’ ^14,15^.

The V_1_ and V_O_ regions of V-ATPase can undergo reversible dissociation and reassembly to regulate enzyme activity in response to changes in cellular conditions ^16,17^. Glucose depletion in the growth medium of the yeast *Saccharomyces cerevisiae* causes V_1_ and V_O_ to separate, with the V_1_ and V_O_ complexes adopting inhibited conformations and subsequent dissociation of subunit C from the V_1_ complex ^18,19^. On restoration of glucose to the growth medium, the enzyme complex known as RAVE (<u>r</u>egulator of the <u>A</u>TPase of <u>v</u>acuoles and <u>e</u>ndosomes) mediates V-ATPase reassembly, likely by bringing V_1_ and V_O_ into close proximity and inducing conformational changes needed for their interaction ^20–22^. When separated from the V_1_ region, the V_O_ complex adopts rotational State 3 ^12^, with the isolated intact V_1_ complex adopting rotational State 2 but sampling all three rotational states after separation of subunit C ^19^.

The initial assembly of the V_O_ complex in the endoplasmic reticulum (ER) membrane of yeast requires the assembly factors Vma12p, Vma21p, and Vma22p ^23–25^. These proteins, which are homologous with mammalian TMEM199, VMA21, and CCDC115, respectively, are necessary for V-ATPase activity in cells but are not found in the mature enzyme. The mammalian proteins are required for functional lysosomes ^26,27^, with their deficiency associated with X-linked myopathy with excessive autophagy ^27^, follicular lymphoma ^28^, and congenital disorders of glycosylation ^29–32^. Vma21p is an integral membrane protein that participates in V_O_ assembly but also accompanies fully assembled V_O_ out of the ER ^24,33^. Following assembly of V_1_ with V_O_, a C-terminal ER retrieval motif on Vma21p allows it to be transported back to the ER to participate in additional rounds of V_O_ assembly ^24,33^. Vma12p, which contains a transmembrane region, forms a complex with the soluble protein Vma22p (the Vma12-22p complex) that stabilizes subunit a and mediates its assembly with the c ring ^23,25,34^.

To understand the structural basis by which Vma21p and the Vma12-22p complex facilitate V_O_ assembly, we used cryoEM to determine structures of yeast V_O_ assembly intermediates bound to both species. The structures show that the Vma12-22p complex binds both a partially assembled V_O_ complex that lacks subunits a, e, and f, as well as fully assembled V_O_ with its c ring in a different rotational state than found in mature V_O_. In contrast, Vma21p binds complexes that include mature V_O_, the assembly intermediate that contains Vma12-22p, and a small population of particles that appear to consist of loosely packed c rings bound to the uncharacterized proteins YAR027W and YAR028W. Together, these structures suggest how Vma12-22p and Vma21p carry out their functions, and the sequence of events in V_O_ assembly.

## Results

### Vma12-22p forms complexes with V_O_ and V_O_ lacking subunits a, e, and f

To understand how the Vma12-22p complex facilitates V_O_ assembly, we integrated DNA encoding a 3×FLAG tag into the yeast chromosome 3’ of the gene for Vma12p. Following yeast growth, cell membranes were isolated, solubilized with detergent, and protein was purified by anti-FLAG affinity chromatography. The resulting protein preparation was subjected to structural analysis by cryoEM (Fig. S1A). This preparation gave rise to two distinct 3D structures. The first structure resembles the V_O_ complex but with additional proteins bound and has a nominal overall resolution of 2.6 Å, with local resolutions ranging from 2.3 to 4.8 Å (Fig. 1A, Fig. S1B *blue curve*, Fig. S1C and D). The map enabled construction of an atomic model (Fig. 1B, Table S1), which included all the known yeast V_O_ subunits and subunit F from the V_1_ region. The map also allowed fitting and refinement of models of Vma12p and Vma22p from AlphaFold ^35^ (Fig. 1A and B, *yellow* and *blue* densities and models). Consequently, this complex is designated the V_O_:Vma12-22p complex. The second structure, which was obtained from ~27% of the particle images, appears as a partial V_O_ complex lacking subunits a, e, and f but with Vma12-22p bound (Fig. 1C). This map was refined to 3.1 Å resolution, with local resolutions ranging from 2.6 to 6.1 Å (Fig. S1B, *green curve*, Fig. S1E and F). The map enabled construction of an atomic model consisting of the c ring (subunits c_8_, c’, c”, and Voa1p), subunit d, the Vma12-22p complex, and subunit F (Fig. 1D, Table S1). Based on its subunit composition, we designate this assembly the V_O_Δaef:Vma12-22p complex.

**Figure 1.**
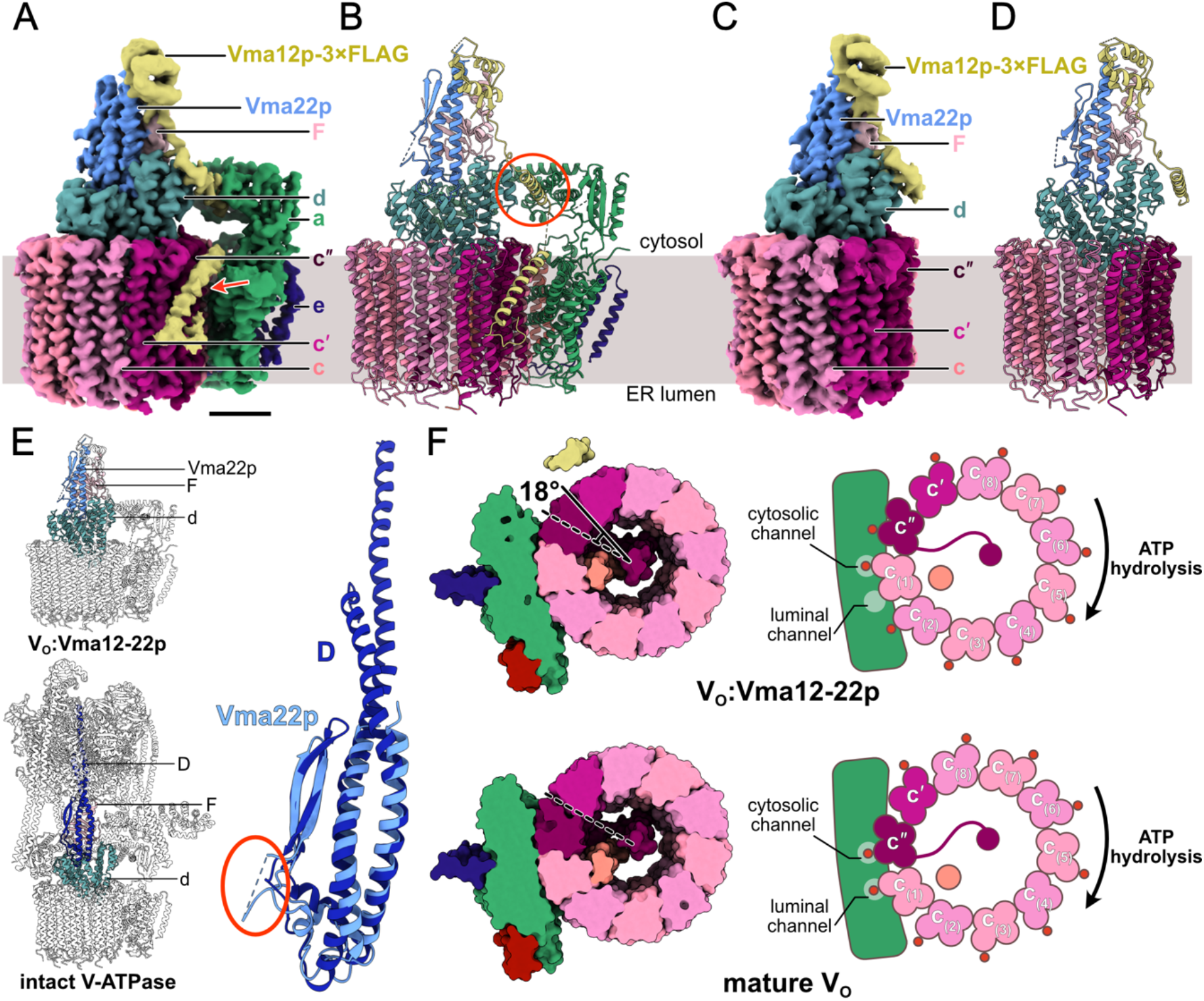
Structure of the V_O_:Vma12-22p complex. **A,** CryoEM map of the V_O_:Vma12-22p complex. Scale bar, 25 Å. **B,** Atomic model for the V_O_:Vma12-22p complex. **C,** CryoEM map of the V_O_Δaef:Vma12-22p complex. **D,** Atomic model for the V_O_Δaef:Vma12-22p complex. **E,** Structural comparison of Vma22p and subunit D from the V_1_ region of V-ATPase. **F,** Comparison between cross-sections through the V_O_:Vma12-22p complex and the mature V_O_ complex (PDB: 6O7T) shows the difference in c ring rotational state.

Interaction between Vma12-22p complex and subunit a is known to be transient ^34^. However, we were still concerned that the presence of V_o_Δaef:Vma12-22p in the dataset could be due to dissociation of subunit a, e, and f from V_O_:Vma12-22p during cryoEM specimen preparation, rather than representing a species that exists in the ER membrane. A structure of V_O_Δaef has not been described in previous cryoEM analysis of V_O_ complexes ^12,13,36,37^ but it was not clear whether these broken complexes were removed during image analysis in those studies. Therefore, we reprocessed a published dataset of cryoEM images of V_O_ complex ^36^ that had been purified and frozen in conditions equivalent to those used for the Vma12p-3×FLAG preparation. The results showed only intact V_O_ (Fig. S2A), indicating that subunits a, e, and f are unlikely to separate from V_O_ during grid freezing, and suggesting that the V_O_Δaef:Vma12-22p complex is not an artifact of cryoEM specimen preparation. However, we cannot exclude the possibility that the interaction between subunit a and the c ring in V_O_:Vma12-22p is unstable, causing subunits a, e, and f to dissociate in the purified complex, although previous work has suggested that Vma12p and Vma22p stabilize the interaction of subunit a with the c ring ^34^.

### Vma22p mimics subunit D from V_1_ and blocks V_1_ binding

In both the V_O_:Vma12-22p and V_O_Δaef:Vma12-22p complexes, Vma22p sits on top of subunit d in a site that is occupied by subunit D from the V_1_ region in intact V-ATPase ^14,38^ (Fig. 1E, *left*). Vma22p consists primarily of a twisted pair of a helices abutted by two antiparallel β strands (Fig. 1E *right, light blue*). This fold resembles the fold of subunit D (Fig. 1E *right, dark blue*), which shares 18 % amino acid sequence similarity and 10 % amino acid sequence identity with Vma22p (Fig. S2B). The a helical structure in Vma22p is less than half the length of the corresponding a helical region in subunit D, making Vma22p too short to serve the same role as subunit D in the V_1_ complex. Thus, the structure of Vma22p suggests that it would block V_1_ from binding subunit d, which would prevent premature attachment of V_1_ to partially assembled V_O_ complexes in the ER membrane. Blocking V_1_ from binding V_O_ would prevent proton pumping into the ER lumen, which is maintained at near neutral pH ^39,40^. Compared to subunit D, Vma22p possesses an additional disordered loop of unknown function comprising 35 residues C-terminal to its two β strands (Fig. 1E *right, circled in red*).

Surprisingly, V-ATPase subunit F is found interacting with Vma12-22p (Fig. 1A-D, *light pink*). Subunit F is a small protein that normally binds subunit D in both intact V-ATPase and the dissociated V_1_ complex, but is not part of the dissociated V_O_ complex. The role of subunit F when bound to subunit D is not known. However, both the V_O_:Vma12-22p and V_O_Δaef:Vma12-22p structures show the N-terminal domain of subunit F in contact with Vma22p while its C-terminal a helix interacts with both Vma12p and Vma22p (Fig. S2C). This interaction with both Vma12p and Vma22p suggests that subunit F may help mediate association of the two proteins. Deletion of the gene for subunit F leads to reduced levels of V_O_ subunits a and c in yeast, with normal levels of subunit d but reduced association of subunit d with the vacuolar membrane ^41^. This phenotype is consistent with the association between subunit F and Vma12-22p being necessary for function of the assembly factors.

### Vma12p helps attach subunit a to the c ring

Vma12p possesses a compact folded N-terminal domain comprising three a helices and two β strands (Fig. 1A-D, *yellow*). In both the V_O_:Vma12-22p complex and the V_O_Δaef:Vma12-22p complex, this N-terminal domain interacts with subunit F and Vma22p. Following the N-terminal domain of Vma12p in the V_O_:Vma12-22p structure, there is an extended linker and an a helix that contacts both a loop (residues 48-53) from subunit a and two short a helices (residues 27-35 and 39-48) from subunit d (Fig. 1B, *red circle*, Fig. S2D). This a helix from Vma12p is followed by a second extended linker and a membrane-embedded a helix that is tilted relative to the plane of the lipid bilayer (Fig. 1A, *red arrow*). Vma12p is predicted to have two C-terminal transmembrane a helices, with its C terminus on the cytosolic side of the membrane ^25,42^. However, apparent flexibility in this region of the protein prevents the second transmembrane a helix from being resolved in the cryoEM map. Neither of the two transmembrane a helices from Vma12p could be resolved in the V_O_Δaef:Vma12-22p map (Fig. 1C). To ensure that flexibility in the C-terminal transmembrane a helices from Vma12p is not due to the C-terminal 3×FLAG tag introduced in the protein, we constructed a second yeast strain with the affinity tag at the C terminus of Vma22p instead. CryoEM of protein purified from this second strain resulted in maps of V_O_:Vma12-22p and V_O_Δaef:Vma12-22p that are indistinguishable from the maps obtained from the Vma12p-3×FLAG preparation (Fig. S2E and F). This finding indicates that flexibility in the C-terminal transmembrane a helices of Vma12p is not due to the incorporation of the affinity tag. The equivalence of the two structures obtained from the Vma12p-3×FLAG and Vma22p-3×FLAG preparations also shows that Vma12p and Vma22p form an obligate heterodimer. It appears that interaction between Vma12p and the N-terminal domain of subunit a reduces mobility in one of the transmembrane a helices from Vma12p, allowing it to be resolved in the cryoEM map. This transmembrane a helix from Vma12p does not interact directly with the transmembrane C-terminal domain of subunit a or the c ring. Therefore, other than anchoring Vma12-22p to the membrane, the role of the tilted transmembrane domain in Vma12p is unclear. In contrast, the interaction between subunit a, subunit d, and the short a helix N-terminal of the transmembrane domain in Vma12p is consistent with Vma12-22p’s proposed role of stabilizing subunit a and recruiting it to V_O_^34^.

### The c ring rotation differs between V_O_:Vma12-22p and mature V_O_ complex

The c ring has an asymmetric distribution of proton-carrying Glu residues, with the protonatable Glu residue of subunit c” found on the second ring-forming a helix, while the protonatable Glu residues of the eight c subunits and one c’ subunit are located on their fourth ring-forming a helices ^12^ (Fig. 1F, *right*). Previous structures of yeast V_O_ found the complex in rotational State 3 with Glu108 from subunit c” aligned with the cytosolic half channel ^12,13,36,37^, while Glu137 from subunit c(1) is aligned with the luminal half channel (Fig. 1F, *lower*). Compared to these structures of mature V_O_, the V_O_:Vma12-22p complex shows an ~18° rotation of the c ring in the ATP-hydrolysis direction (Fig. 1F, *upper*). As a result, in the V_O_:Vma12-22p complex, Glu137 from subunit c(1) is aligned with the cytosolic half channel and Glu137 from subunit c(2) is far from the luminal half channel (Fig. 1F, *upper right*). A previous study of yeast V_O_ complexes by cryoEM detected a similar ~14° rotation of the ring in a 3D class from ~5% of particle images ^43^. Both the small population of particle images in the class and molecular dynamic simulations indicated that this conformation is less energetically favourable than V_O_ in rotational State 3 ^43^. Comparison of the V_O_:Vma12-22p and mature V_O_ structures shows that even with the ~18° rotation of the c ring relative to the C-terminal domain of subunit a, the interface between subunit d and N-terminal domain of subunit a remains unchanged (Fig. S2G i), as shown previously ^43^. Further, rotation of the c ring from its V_O_:Vma12-22p conformation to the mature V_O_ conformation would cause a clash between Vma12p and subunit a that would prevent Vma12-22p from binding to V_O_ (Fig. S2G ii, *orange circle*). Therefore, the structure suggests that after Vma12p recruits subunits a, e, and f to the csc’c”dVoa1p ring, relaxation of the c ring into its energetically favoured conformation disrupts the interaction between Vma12-22p and V_O_, releasing Vma12-22p and resulting in the mature V_O_ structure. However, this release of Vma12-22p from V_O_ must occur either slowly or with other proteins involved, or else it would not be possible to isolate the V_O_:Vma12-22p complex biochemically.

### Vma21p binds the V_O_ complex in multiple positions around the c ring

To understand how the activity of Vma21p relates to the V_O_ assembly process, we next investigated V_O_ complexes that contain Vma21p. As with Vma12-22p, we constructed a yeast strain with DNA for a 3×FLAG tag integrated in the chromosome 3’ of the *VMA21* gene. When grown on medium containing zinc, the Vma21p-3×FLAG strain exhibited growth comparable to wildtype yeast, while a ΔVma21p strain showed the expected *VMA*^-^ V-ATPase deficiency phenotype (Fig. S3), indicating that despite Vma21p’s C-terminal Lys-Lys-Glu-Asp ER-retrieval motif ^24^ the C-terminal 3×FLAG tag does not prevent assembly of functional V-ATPase complexes.

CryoEM of protein purified from the Vma21p-3×FLAG strain led to a 3D map resembling the mature V_O_ complex in rotational State 3 that we designate the V_O_:Vma21p complex (Fig. 2A, Fig. S4A). The nominal overall resolution of the map is 3.1 Å with local resolutions ranging from 2.2 to 7.5 Å (Fig. S4B, *orange curve*, Fig. S4C and D). The map enabled construction of an atomic model through a combination of rigid body fitting and refinement of a published V_O_ model ^36^ and an AlphaFold predicted model for Vma21p ^35^ (Fig. 2B, Table S1). Clearly defined density accommodated a single copy of Vma21p (Fig. 2A and B, *purple*). No conformational differences could be detected between the V_O_ part of V_O_:Vma21p and a mature V_O_ structure (Fig. S5A). In the structure, Vma21p forms a transmembrane a helical hairpin wedged between two adjacent c subunits near the luminal half channel of subunit a. The N-terminal a helix from Vma21p interacts with the second a helix of subunit c_(2)_ and the fourth a helix of subunit c_(3)_, while the C-terminal a helix from Vma21p contacts the second and fourth a helices of subunit c_(2)_ (Fig. S5B).

**Figure 2.**
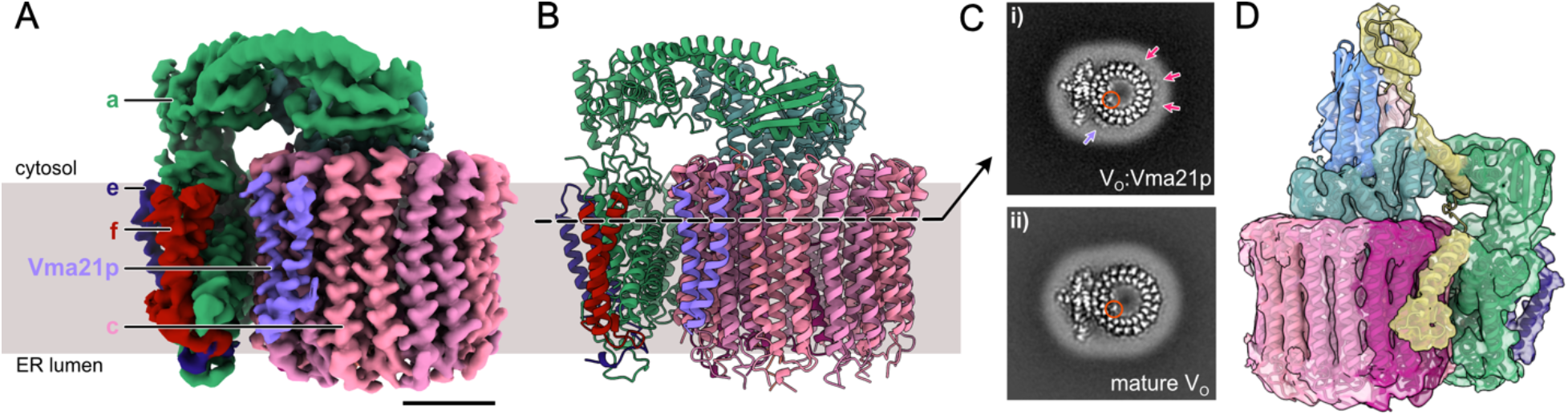
Structure of the V_O_:Vma21p complex. **A,** CryoEM map of the V_O_:Vma21p complex. Scale bar, 25 Å. **B,** Atomic model for the V_O_:Vma21p complex. **C,** A slice through the V_O_:Vma21p map (i) showing additional protein densities surrounding the c ring compared to the mature V_O_ complex (EMDB: 0644) (ii). Purple arrow, strong density for Vma21p. Magenta arrows, weaker density for Vma21p. Orange circle, density for Voa1p. **D,** A subset of the particle images from the Vma21p-3×FLAG dataset gave a map of V_O_:Vma12-22p complex, shown here with the atomic model of V_O_:Vma12-22p from the Vma12p-3×FLAG dataset fitted.

The well-defined single Vma21p binding site in V_O_:Vma21p involves only two of the eight c subunits and does not appear to be unique within the c ring. To understand Vma21p’s apparent preference for a single binding site, cross-sections through the V_O_:Vma21p map were inspected and compared to cross-sections through a map of mature V_O_ complex (Fig. 2C). The two a helices from the well-resolved Vma21p are clear in the cross-section (Fig. 2C i, *purple arrow*). However, features corresponding to additional similar but lower density a helices can be seen surrounding the c ring (Fig. 2C i, *magenta arrows*). These additional densities, which are not seen in a map of the mature V_O_ complex determined by equivalent methods ^36^ (Fig. 2C ii), suggest that multiple copies of Vma21p bind around the c ring but with a low binding occupancy in all but one site. In its preferred site, Vma21p does not interact with the nearby subunit a and it is not clear why this site has a high occupancy while the others sites have a low occupancy. It is worth noting that in the V_O_:Vma12-22p complex this Vma21p would be in contact with subunit a. However, the V_O_:Vma12-22p map does not show density for Vma21p at this position. Cross sections through the map also show density for the C-terminal transmembrane a helix from Voa1p (Fig. 2C i, *orange circle*), which co-precipitates with Vma21p ^44^ and can be seen in maps of V_O_Δaef:Vma12-22p and V_O_:Vma12-22p (Fig. 1F, *orange*), as well as mature V_O_ complex (Fig. 2C ii, *orange circle*) ^12,13,36^.

The V_O_:Vma21p structure appears to show the assembled mature V_O_ complex that is escorted from the ER by Vma21p. A Vma21p mutant with a defective ER-retrieval motif was previously found to interact with intact V-ATPase ^33^ and V_1_ subunits were detected in mass spectrometry of the Vma21p-3×FLAG protein preparation (Table S2). However, we did not identify particle images from the Vma21p-3×FLAG dataset that show intact V-ATPase, suggesting any interaction between Vma21p and intact V-ATPase is unstable.

### Vma21p binds a V_O_:Vma12-22p complex and a complex containing YAR027W and YAR028W

3D classification of the cryoEM dataset from Vma21p-3×FLAG tagged protein identified two additional populations of particle images. The first allowed calculation of a 3D map at 4.4 Å resolution that was indistinguishable from the V_O_:Vma12-22p complex (Fig. 2D, Fig. S4B, *green curve*, Fig. S4E and F). Density for Vma21p could not be detected in this map, suggesting that again Vma21p interacts with the c ring, but with variable positions that leads to incoherent averaging in 3D reconstruction.

The second unique class of particle images isolated with 3×FLAG tagged Vma21p allowed calculation of a 3D map that resembles a c ring with additional protein densities attached (Fig. 3A). Only 19,849 out of 607,746 particle images from the dataset contribute to this structure, leading to a poor overall resolution of 5.7 Å, with local resolutions between 4.6 and 9.2 Å (Fig. S4B, *pink curve*, Fig. S4G and H). To identify the additional protein density seen in this map, we inspected a list of proteins identified in the preparation by mass spectrometry of tryptic fragments (Table S2) and attempted to fit AlphaFold predicted structures of candidate proteins into the map. This process identified two candidate proteins, YAR027W and YAR028W. The amino acid sequences for these two putative integral membrane proteins are 58% identical and 73% similar (Fig. S6), with AlphaFold predicting nearly indistinguishable folds for the two proteins (Fig. 3B). Due to the low resolution of the cryoEM map and the similarity in the predicted folds of YAR027W and YAR028W, the two proteins could not be distinguished in the map (Fig. 3C). A view of the map perpendicular to the plane of the detergent micelle shows nine copies of YAR027W or YAR028W surrounding a ring that appears to consist of nine loosely packed c subunits (Fig. 3D, *dashed circles*). Therefore, we designated this assembly the YAR027W/028W:c9 complex. Due to the limited resolution of the map, the c, c’, and c” subunits could not be distinguished in the ring. Examination of the fold and surface properties of YAR027W and YAR028W suggests that they each form two N-terminal transmembrane a helices that are buried in the detergent micelle, with a C-terminal soluble domain likely on the cytosolic side of the ring (Fig. 3E, *blue* and *yellow* surfaces).

**Figure 3.**
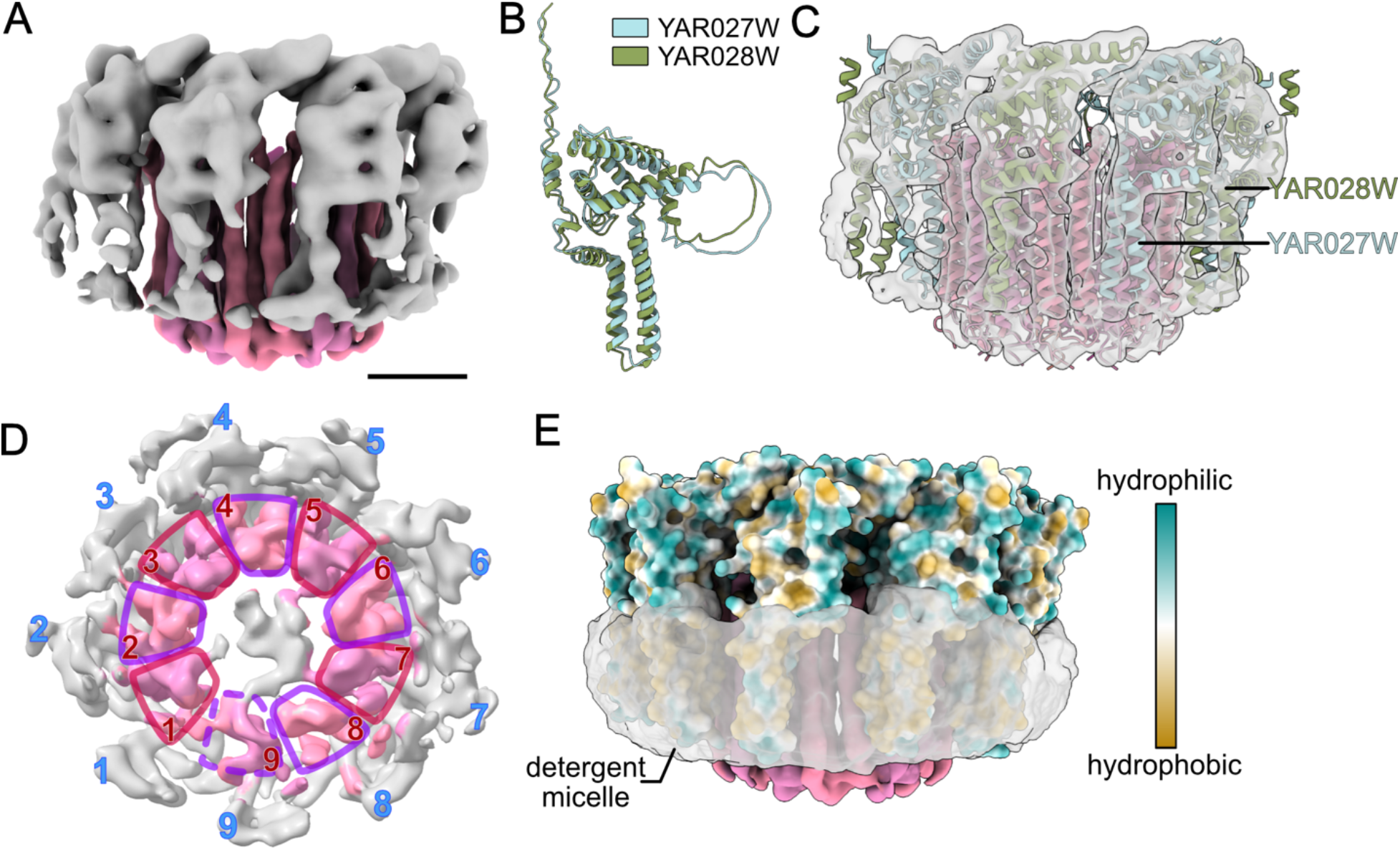
Structure of YAR027W and YAR028W binding to a loosely packed c ring. **A,** A subset of the particle images from the Vma21p-3×FLAG dataset produced a map that resembles a c ring with additional protein densities attached. Scale bar, 25 Å. **B,** Comparison of the AlphaFold predicted structures of YAR027W and YAR028W. **C,** Fitting of AlphaFold predicted structures of YAR027W, YAR028W, and c subunits into the map.**D,** A view of the map perpendicular to the plane of the detergent micelle shows nine copies of YAR027W or YAR028W surrounding a structure that resembles nine loosely packed c ring protomers. **E,** Surface hydrophobicity of YAR027W and YAR028W compared with the position of the detergent micelle in the map.

The *YAR027W* and *YAR028W* genes are members of the *S. cerevisiae DUP240* multigene family, which contains non-essential membrane-associated proteins ^45,46^. The two proteins have been shown in high-throughput studies to interact with each other, as well as multiple other proteins ^47,48^ but have not, to our knowledge, been shown to interact with components of V-ATPase. YAR027W, also known as UIP3, was reported to localize to the nuclear envelop ^46^. A recent study showed that yeast cells that were resistant to killer toxin K28 became sensitive to the toxin when the gene *YAR028W* was deleted ^49^. While it is tempting to speculate that YAR027W and YAR028W are somehow involved in c ring assembly, the resolution of the map precludes determining the precise locations of YAR027W, YAR028W, c, c’, or c” in the complex. Numerous questions related to the interaction between YAR027W and YAR028W require investigation. For example, it is unknown if the two highly similar proteins are functionally redundant and if a double knockout compromises V-ATPase assembly. It also remains unknown if most of the YAR027W and YAR028W in the cell is bound to V-ATPase c rings with Vma21p, or if this population is a minor portion of the YAR027W and YAR028W. Further, it is unclear if YAR027W and YAR028W interact with other proteins beside the components of the c ring and Vma21p, and if these interactions are somehow related to their interaction with V-ATPase. A high-resolution structure of YAR027W, YAR028W, the c ring, and Vma21p may help answer some of these questions. While extensive further study is clearly required, the results presented here indicate that Vma21p interacts with a complex that involves c subunits, YAR027W, and YAR028W.

The C-terminal KKED ER-retrieval motif of Vma21p allows the protein to travel back to the ER membrane after escorting the fully assembled V_O_ complex out of the ER. Mutating the motif to QQER results in Vma21p’s mis-localization to the vacuolar membrane ^24^. A recent study showed that removal of this retrieval motif on human VMA21 results in the protein becoming enriched in lysosomes, eventually leading to activation of autophagy ^28^. The structures obtained from Vma21p-3×FLAG purification suggests that Vma21p binds all the c ring containing V_O_ subcomplexes. By binding to these complexes, the ER-retrieval motif on Vma21p would return any partially assembled V_O_ complexes that are transported to the Golgi back to the ER membrane. Therefore, removal of Vma21p is required for the mature V_O_ complex to remain in the Golgi or travel further to its designated cellular location.

## Discussion

Unsurprisingly, owing to their importance for V_O_ assembly, mutations in the human homologues of Vma12p, Vma22p, and Vma21p have been linked to disease. The results presented here provide a structural context to understand these mutations. Mutations of the human homologue of Vma22p, CCDC115, are found in patients affected by congenital disorders of glycosylation ^30,31^. The human mutation D11Y maps onto a region at the N terminus of Vma22p that is not resolved in the structure. However, L31S in CCDC115 maps onto Leu34 in the yeast structure, where mutation could affect the binding of subunit F (Fig. S7A, *blue circle with cyan fill*). Mutations in TMEM199, the human homologue of Vma12p, are also found in the same disease ^32^ and map to the N-terminal domain of Vma12p ^32^ (Fig. S7A, *yellow circles with cyan fill*). These residues do not appear to be involved in protein-protein interactions, but their mutation may disrupt the fold of Vma12p. Mutations of the human gene *VMA21* have been reported in X-linked myopathy with excessive autophagy ^27^ and follicular lymphoma ^28^. Three of the reported mutations (equivalent to Thr35, Pro36, and Thr38 in the yeast Vma21p), are found in the luminal loop of the protein (Fig. S7B). The remaining reported mutations map to the transmembrane region of yeast Vma21p where it interacts with c subunits and could disrupt this interaction ^28,29^ (Fig. S7B).

The structures described above suggest the sequence of events that occur during V_O_ assembly in the ER membrane and subsequent binding of V_1_ in the Golgi (Video. S1, Fig. 4). The structure consisting of a partial c ring with YAR027W and YAR028W purified via Vma21p-3×FLAG may or may not be on the assembly pathway and requires further investigation. The c ring assembles around Voa1p ^13,36^, with binding of subunit d to the c_8_c’c”Voa1p ring ^33^ masking the ER-retrieval motif of Voa1p ^13^. The V_O_Δaef:Vma12-22p structure indicates that the Vma12-22p complex binds this fully assembled ring prior to interaction with subunits a, e, and f (Fig. 4A). Vma12-22p helps recruit and secure the interaction of subunit a with the c ring ^34^ through interaction of Vma12p with subunits a and d (Fig. 4B). During recruitment of subunits a, e, and f, Vma22p mediates the interaction between Vma12p and subunit d and prevents premature attachment of the V_1_ complex to the partially assembled V_O_ complex. Subunit F does not directly participate in the assembly process but mediates the association of Vma12p and Vma22p. The fully assembled V_O_ complex with Vma12-22p can then transition to the energetically favored conformation of the c ring, disrupting the Vma12-22p binding site on V_O_ and removing Vma12-22p from the complex (Fig. 4C). This release of Vma12-22p is required for the subsequent assembly of V_1_ with V_O_. The structures presented here suggest that Vma21p binds all assembly intermediates that contain a c ring, with the ER-retrieval motif from Vma21p ensuring that partially-assembled complexes are returned to the ER membrane (Fig. 4A-C).

**Figure 4.**
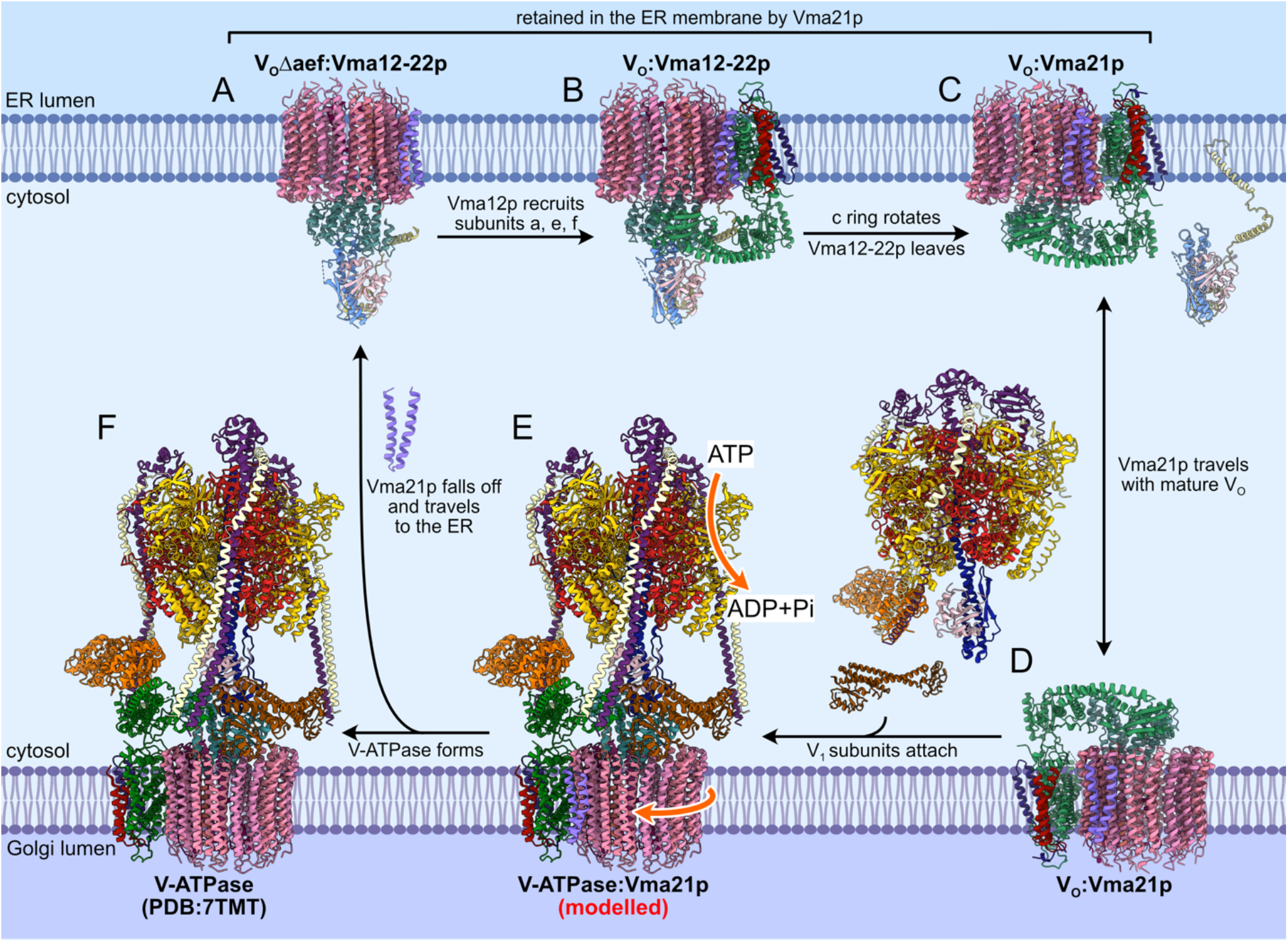
Sequence of events in V_O_ assembly. **A,** Vma12-22p binds a complex containing the c ring, subunit d, and Vma21p. **B,** Vma12p recruits subunits a, e, and f to the rest of V_O_. **C,** V_O_ relaxes to the preferred orientation of the c ring, releasing Vma12-22p. Vma21p ensures V_O_ assembly intermediates are retrieved to the ER membrane. **D,** Vma21p accompanies the fully assembled V_O_ to the Golgi membrane. **E,** V_1_ΔC and subunit C bind V_O_ and initiate rotary catalysis. **F,** Rotation of the c ring releases Vma21p, which travels back to the ER membrane, leaving functional V-ATPase in the Golgi membrane. Please see Video 1.

Despite the ER-retrieval motif of Vma21p, the assembly factor is found in a subpopulation of V_O_ complexes on COPII-coated transport vesicles ^33^, which facilitate V_O_ transport from the ER membrane to the Golgi membrane (Fig. 4D). In the absence of Vma12-22p absent, this V_O_:Vma21p complex is capable of interacting with V_1_, presumably with the RAVE complex mediating binding of V_O_, V_1_ΔC, and subunit C ^19,22^ (Fig. 4E). The attachment of V_1_ would initiate rotary catalysis, so that rotation of the c ring drives dissociation of Vma21p (Fig. 4F). The removal of Vma21p allows intact V-ATPase to remain in the Golgi membrane or travel further to the vacuole, while free Vma21p returns to the ER membrane owing to its ER-retrieval motif. This substitution of Vma21p with V_1_ is consistent with simultaneous loss of Vma21p and binding of V_1_ to V_O_ observed previously ^33^.

Overall, the data presented here provide a structural context for understanding disease-causing mutations in CCDC115, TMEM199, and VMA21. Furthermore, the structures suggest how Vma12-22p and Vma21p coordinate their known roles in the assembly of the V_O_ complex and formation of an active intact V-ATPase after departure of V_O_ from the ER membrane. They also reveal an unexpected protein complex involving the V-ATPase c ring, Vma21p, and the uncharacterized proteins YAR027W and YAR028W. Importantly, the structures illustrate how Vma21p and Vma12-22p play central roles in both V-ATPase assembly and quality control.

## Author contributions

HW purified the protein samples and performed cryoEM and associated analysis. SAB prepared the Vma12p-, Vma21p-, and Vma22p-3×FLAG yeast strains and performed proof-of-principle protein purifications. JLR conceived the study and supervised the research. HW and JLR wrote the manuscript and prepared figures.

## Acknowledgements

We thank Samir Benlekbir for assistance with cryoEM data collection, Spencer Freeman and Sergio Grinstein for discussions about ER-retrieval, and Patricia Kane for a critical reading of the manuscript. JLR was supported by the Canada Research Chairs program. This research was supported by Canadian Institutes of Health Research grant PJT166152 (JLR). CryoEM data was collected at the Toronto High-Resolution High-Throughput cryoEM facility, supported by the Canada Foundation for Innovation and Ontario Research Fund. Mass spectrometry data was collected at the Network Biology Collaborative Center at the Lunenfeld-Tanenbaum Research Institute.

## Data availability

Electron microscopy maps are available from the electron microscopy databank with accession codes EMD-27984 to EMD-27988 and atomic models are available from the protein databank with accession codes 8EAS, 8EAT, 8EAU, and 8EAV.

## Methods

### Yeast Strains

Yeast *S. cerevisiae* strains SABY125, SABY129, and SABY130 were prepared by homologous recombination to integrate DNA sequence for a 3×FLAG tag downstream of the *VMA12, VMA21*, and *VMA22* genes in the background strain BJ2168 (*MATa leu2 trp1 ura3-52 prb1-1122 pep4-3 prc1-407gal2*), respectively. Plasmid pJT1 ^38^ was used to amplify DNA sequence encoding the 3 ×FLAG tag followed by a *URA3* marker. The PCR products were then used to transform BJ2168 by standard lithium acetate transformation. Transformed cells were selected by growth on minimal media lacking uracil, and successful transformants were confirmed by PCR. The *Δ* Vma21p yeast strain YSC6273-201936318 (*vma21*::KanMX*MATa his3Δ1 leu2Δ0 met15Δ0*) was purchased from Horizon Discovery.

### Protein Purification

Yeast strains were grown in 11 L yeast extract peptone dextrose (YPD) media (20 g/L peptone, 20 g/L glucose, 10 g/L yeast extract) supplemented with 100 μg/mL ampicillin and 0.02% antifoam in a Microferm fermenter (New Brunswick Scientific) at 30 °C for one day (> 22 h), with aeration of 34 cubic feet per hour and stirring at 300 rpm. All subsequent steps were performed at 4 °C. Cells were harvested by centrifugation at 4000 ×g for 15 min and resuspended in 1 mL/g lysis buffer (8 g/L NaCl, 0.2 g/L KCl, 1.44 g/L Na2HPO4, 0.24 g/L KH2PO4, 80 g/L sucrose, 20 g/L sorbitol, 20 g/L glucose, 5 mM 6-aminocaproic acid, 5 mM benzamidine hydrochloride, 5 mM ethylenediaminetetraacetic acid [EDTA], 10 mg/L phenylmethylsulfonyl fluoride [PMSF], pH 7.4). Cells were lysed with 0.5 mm glass beads (BioSpec) in a bead beater, with six cycles of 1 min of bead beating and 1 min of cooling. Cell debris was removed by centrifugation at 4000 ×g for 15 min. Membranes were then collected by ultracentrifugation (Beckman L-90K, Ti70 rotor) at 145,000 ×g for 40 min, resuspended in 0.5 mL/g lysis buffer using a Dounce homogenizer, and stored at −80 °C prior to protein purification.

Frozen membranes were thawed at room temperature, and all the purification steps were performed at 4 °C. Membranes were solubilized with 1% (w/v) n-dodecyl β-D-maltoside (DDM; Anatrace) and mixed for 30 min. Insoluble material was removed by ultracentrifugation at 158,000 ×g for 1h (Beckman L-90K, Ti70 rotor), and membranes were filtered with a 0.45 μm syringe filter and applied to a 0.5 mL anti-FLAG M2 affinity gel in a column (Millipore Sigma) pre-equilibrated in DTBS (50 mM Tris-HCl, 150 mM NaCl, 0.02% [w/v] DDM, pH 7.4). The column was washed with ten column volumes of DTBS, and protein was eluted with three column volumes of DTBS containing 150 μg/mL of 3×FLAG peptide and one column volumes of DTBS without peptide. Protein was then concentrated to ~200 μL with a 100 kDa molecular weight cutoff (MWCO) Amicon Ultracentrifugal filter (Millipore Sigma) at 1000 ×g, diluted with 4 mL GTBS (50 mM Tris-HCl, 150 mM NaCl, 0.004% [w/v] glyco-diosgenin [GDN; Anatrace], pH 7.4) and concentrated to ~200 μL in the same concentrator. For Vma12p-3×FLAG and Vma22p-3×FLAG, samples were further concentrated to ~1-2 mg/mL with a 100 kDa MWCO Vivaspin 500 centrifugal concentrator (Sartorius) at 12000 ×g. For Vma21p-3×FLAG, the sample was concentrated at 1000 ×g with (Sartorius). Protein concentration was determined by bicinchoninic acid (BCA) assay (Pierce).

### CryoEM Specimen Preparation and Data Collection

Samples were applied to homemade nanofabricated Holey Gold grids with regular arrays of ~2 μm holes ^50,51^. Grids were glow-discharged in air for 2 min, and 1.5 μL of sample was applied to the grids, blotted for 2 s at 4 °C and 80% relative humidity, followed by rapid freezing in liquid ethane with a Leica EM GP2 freezing device. Sample screening was performed with a FEI Tecnai F20 electron microscope operating at 200 kV and equipped with a Gatan K2 Summit direct detector device camera. Images were collected as movies at a magnification of 25000×, with 30 fractions at 5 e^-^/pixel/s and a calibrated pixel size of 1.45 Å. High-resolution data collection was performed with a Titan Krios G3i electron microscope (Thermo Fisher) operating at 300 kV and equipped with a Falcon 4 camera (Vma22p dataset) and a prototype Falcon 4i camera (Vma12p and Vma21p datasets). Automated data collection was performed with the *EPU* software package. Movies consisting of 30 exposure fractions were collected at a nominal magnification of 75000×, corresponding to a calibrated pixel size of 1.03 Å.

### Image Analysis

Movie alignment with patch-based motion correction and estimation of contrast transfer function (CTF) parameters were performed with *cryoSPARC Live*. All other image analysis steps were performed with *cryoSPARC v3* ^52^. After removing movies with undesirable CTF fit, ice thickness, or motion, 4032, 4417 and 4179 movies from the Vma12p, Vma21p, and Vma22p datasets were selected for further processing, respectively. Templates for particle selection were generated from 2D classification of manually selected particle images. Individual particle motion correction was performed ^53^ and datasets were cleaned with multiple rounds of 2D classification and *ab initio* 3D classification. This process provided 692,432 particle images for the Vma12p dataset, 607,746 particle images for the Vma21p dataset, and 462,692 particle images for the Vma22p dataset.

*Ab initio* 3D classification and heterogeneous refinement were applied to all the datasets. For the Vma21p dataset, this procedure allowed identification of classes corresponding to V_O_:Vma21p, V_O_:Vma12-22p, and the YAR027W/028W:c9 ring structure. For both the Vma12p and Vma22p datasets, the process led to identification of V_O_:Vma12-22p and V_O_Δaef:Vma12-22p structures. The classes of V_O_:Vma21p from Vma21p dataset, V_O_:Vma12-22p and V_O_Δaef:Vma12-22p from Vma12p dataset were refined with nonuniform refinement^54^, followed by two rounds of local and global CTF refinement, and nonuniform refinement.

### Atomic model building

The previously published V_O_ model 6O7T ^36^, AlphaFold models of Vma12p, Vma21p and Vma22p ^35^, and subunit F from previously published V-ATPase model 7TMQ ^19^ were used for rigid body fitting into the maps of V_O_:Vma12-22p, V_O_Δaef:Vma12-22p, and V_O_:Vma21p with UCSF Chimera ^55^. Atomic models were constructed by manual model building in *Coot* ^56^, followed by refinement with ISOLDE ^57^, and real space refinement with PHENIX ^58^. Figures were rendered with Chimera ^55^ and UCSF ChimeraX ^59^.

## Supplementary Figures

**Figure S1.**
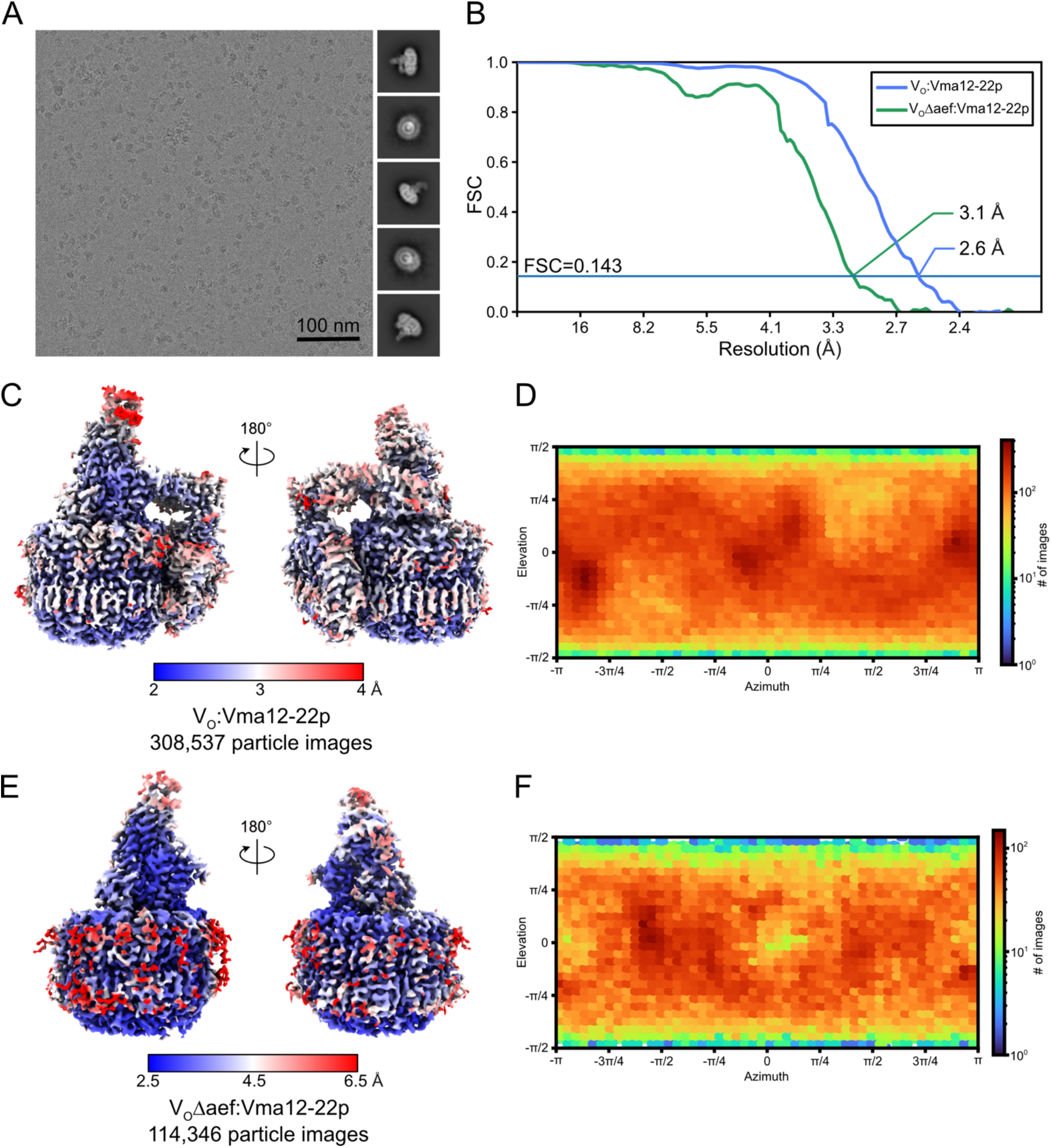
CryoEM of the V_O_:Vma12-22p complex. **A,** Example micrograph and 2D class average images. **B,** Fourier shell correlation curves, corrected for masking, after gold-standard refinement for V_O_:Vma12-22p and V_O_Δaef:Vma12-22p. **C,** Local resolution map for V_O_:Vma12-22p. **D,** Orientation distribution plot for V_O_:Vma12-22p. **E,** Local resolution map for V_O_Δaef:Vma12-22p. **F,** Orientation distribution plot for V_O_Δaef:Vma12-22p.

**Figure S2.**
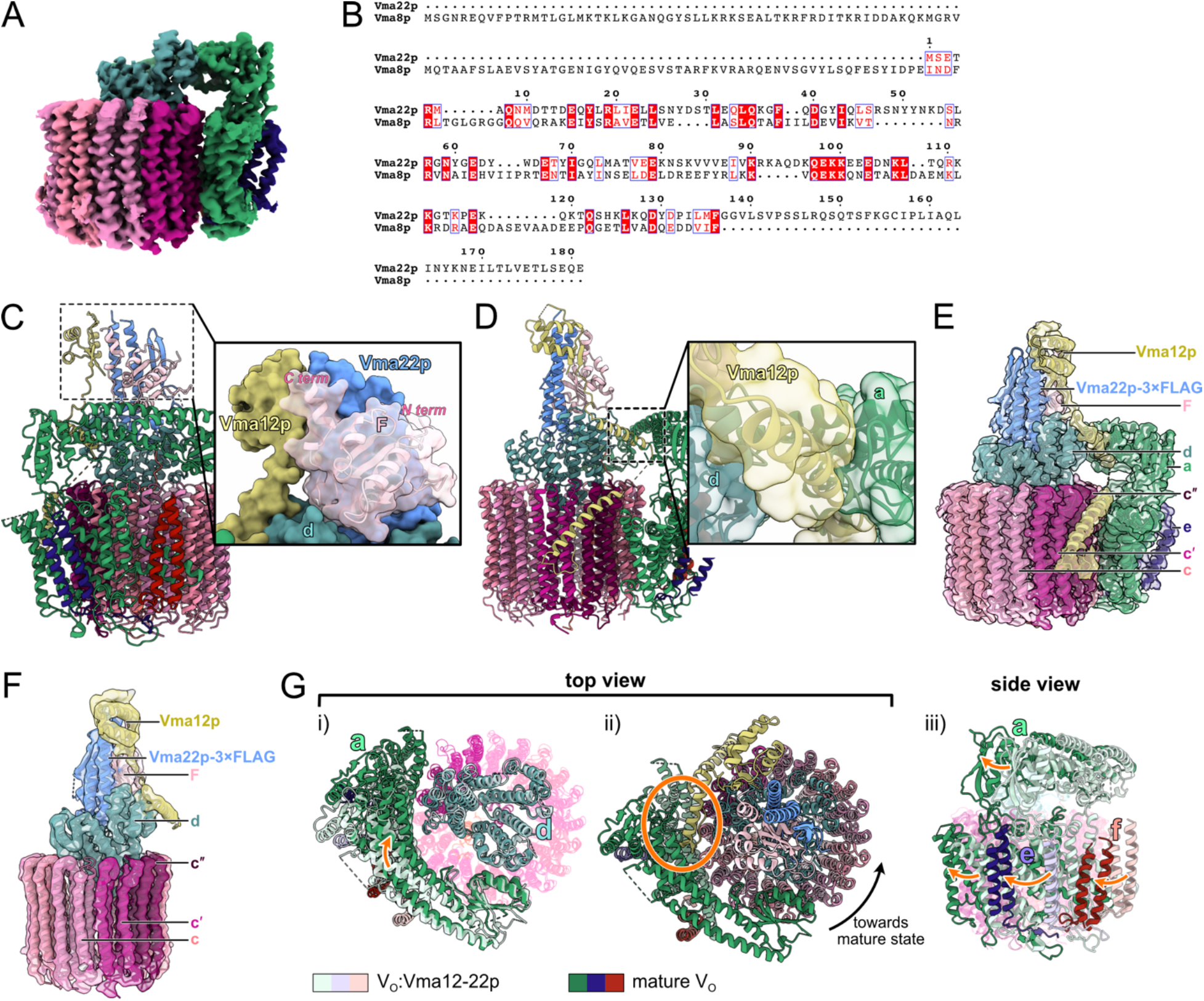
Structural features of the Vma12-22p complex. **A,** Re-analysis of a published data of V_O_ images does not reveal V_O_Δaef complexes. **B,** Protein sequence alignment between Vma22p and Vma8p (subunit D from V_1_ complex). Red boxes indicate identical residues and white boxes indicate similar residues. **C,** Subunit F binds Vma12p and Vma22p and mediates their association. **D,** Vma12p interacts with subunits a and d through an a helix. **E,** CryoEM map of V_O_:Vma12-22p from the Vma22p-3×FLAG preparation with a model of V_O_:Vma12-22p from Vma12p-3×FLAG fitted. **F,** CryoEM map of V_O_Δaef:Vma12-22p from the Vma22p-3×FLAG preparation with a model of V_O_:Vma12-22p from Vma12p-3×FLAG fitted. **G,** Movement of subunits a, e, and f relative to the c ring when the c ring transitions to its orientation in the mature V_O_ complex.

**Figure S3.**
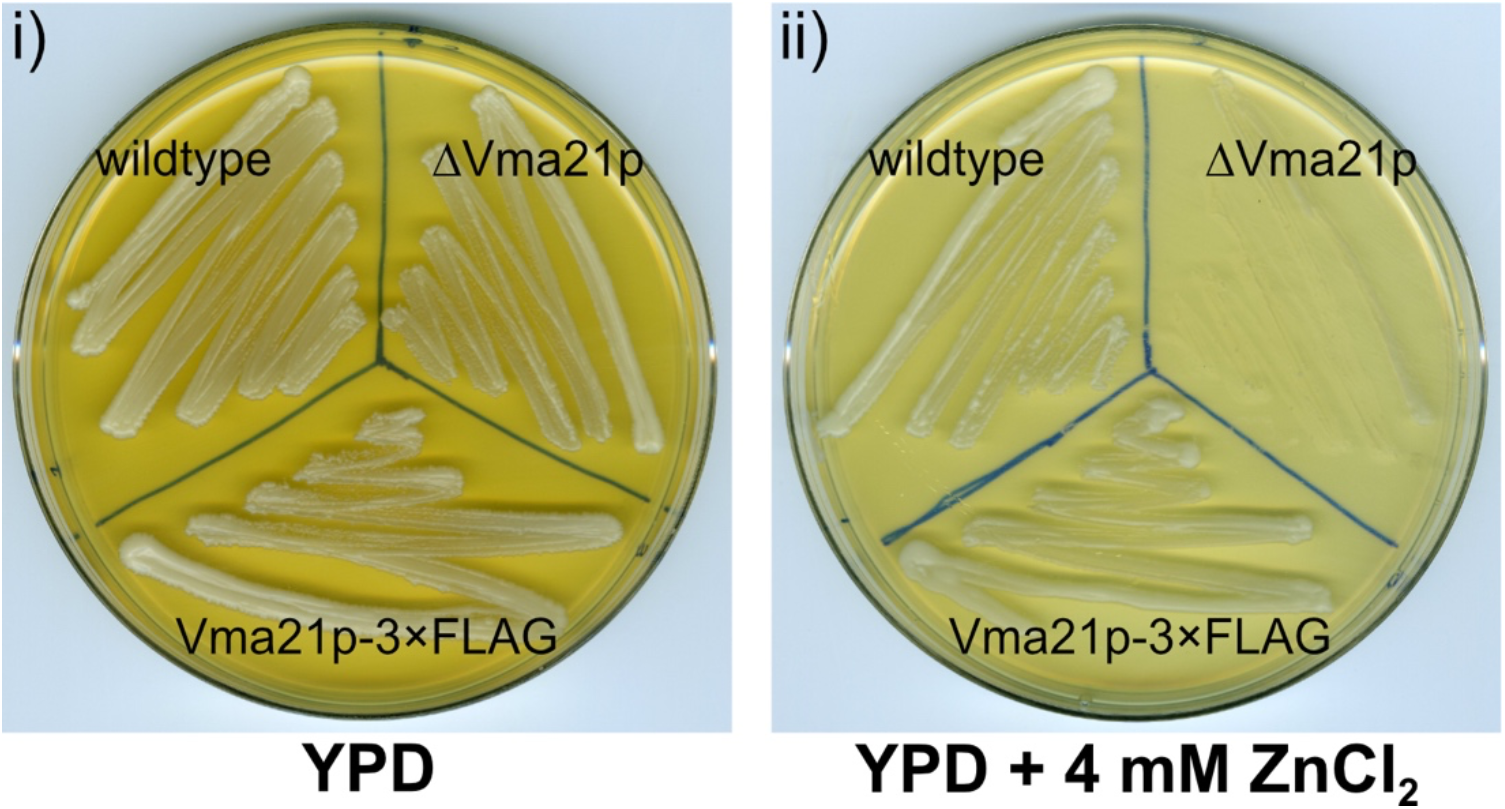
Integration of a 3×FLAG tag to Vma21p does not compromise V-ATPase function. Comparison of yeast growth for wildtype, ΔVma21p, and Vma21p-3×FLAG yeast strains on i) YPD plate, and ii) YPD with 4 mM of ZnCl_2_ plate. Knockout of Vma21p produces a *VMA*^-^ V-ATPase deficiency phenotype but addition of a C-terminal 3×FLAG tag to Vma21p does cause the phenotype.

**Figure S4.**
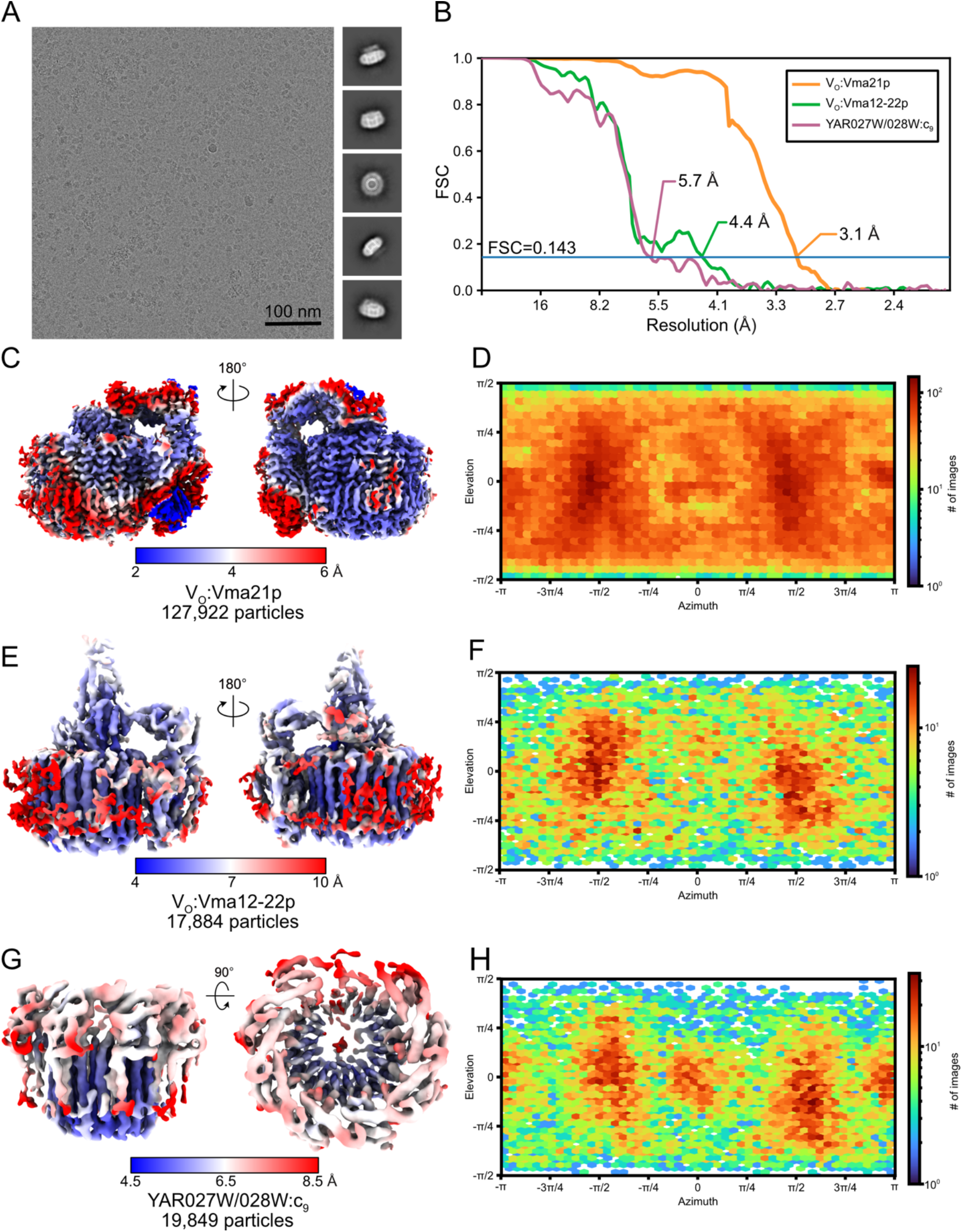
CryoEM of the Vma21p-3×FLAG preparation. **A,** Example micrograph and 2D class average images. **B,** Fourier shell correlation curves, corrected for masking, after gold-standard refinement for V_O_:Vma21p, V_O_:Vma12-22p, and YAR027W/028W:c9. **C,** Local resolution map for V_O_:Vma21p. **D,** Orientation distribution plot for V_O_:Vma21p. **E,** Local resolution map for V_O_:Vma12-22p. **F,** Orientation distribution plot for V_O_:Vma12-22p. **G,** Local resolution maps for YAR027W/028W:c9. **H,** Orientation distribution plot for YAR027W/028W:c9.

**Figure S5.**
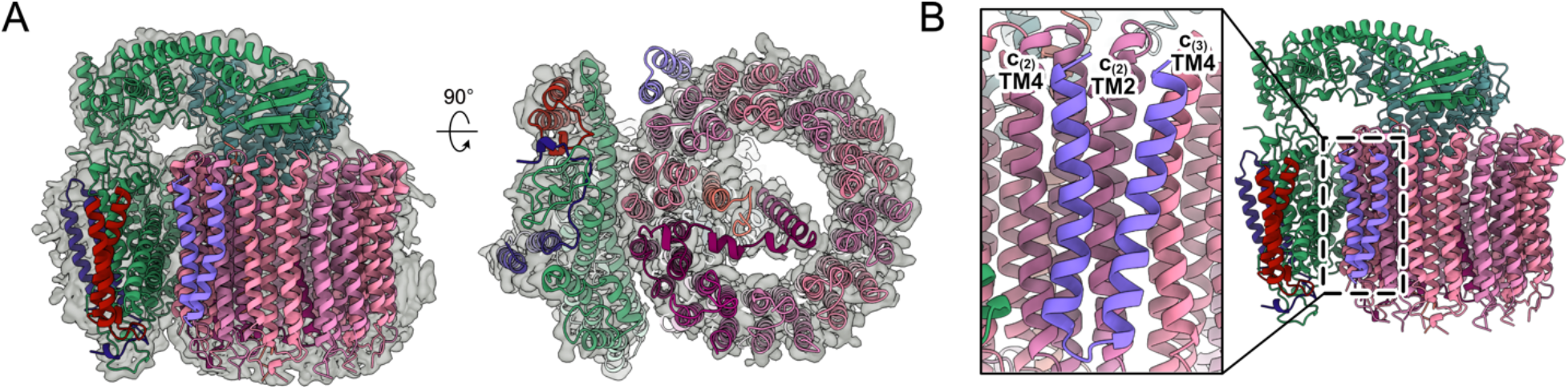
Comparison of the structure of V_O_:Vma21p complex and mature V_O_. **A,** Fitting of the V_O_:Vma21p model into the previously-determined V_O_ map (EMDB: 0644). **B,** Binding location of Vma21p on the c ring.

**Figure S6.**
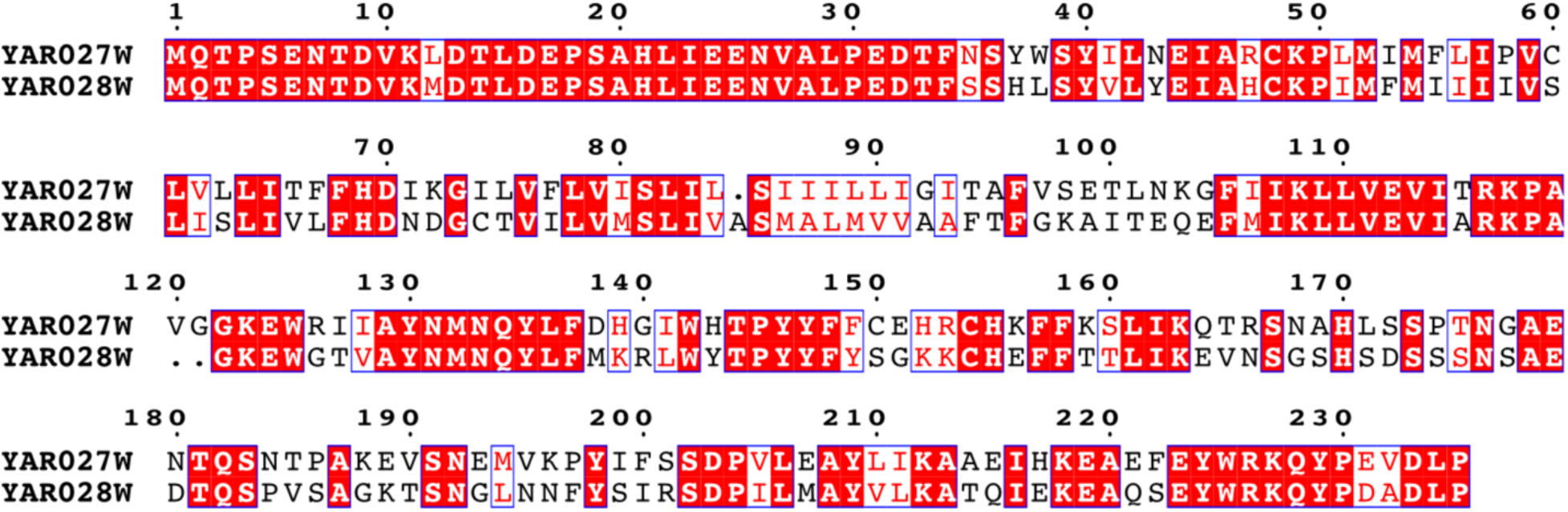
Protein sequence alignment between YAR027W and YAR028W. Red boxes indicate identical residues and white boxes indicate similar residues.

**Figure S7.**
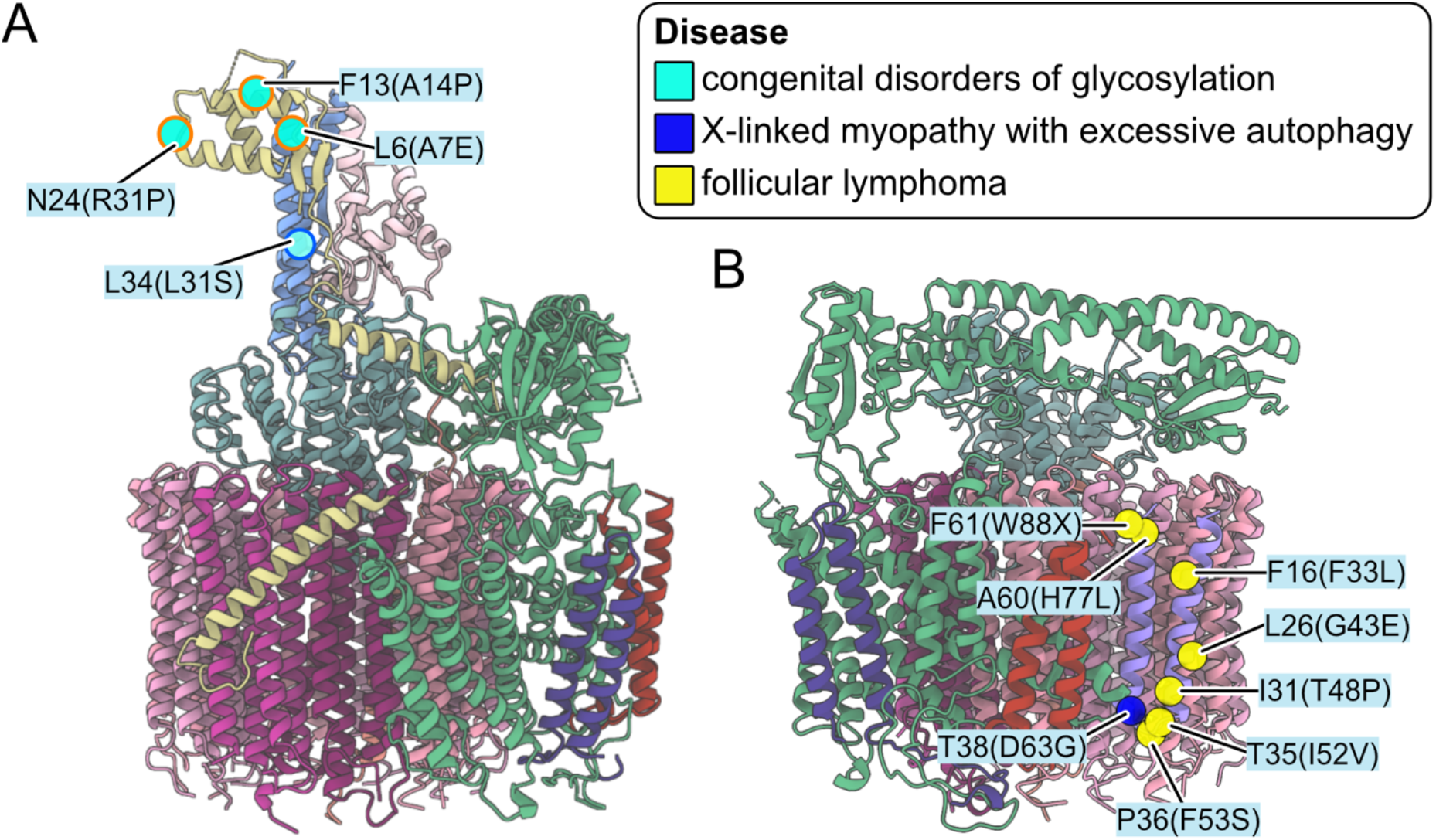
Disease-associated mutations in human TMEM199, VMA21, and CCDC115 that map onto yeast Vma12p, Vma21p, and Vma22p. **A,** Mutations of TMEM199 and CCDC115 associated with congenital disorders of glycosylation that map onto yeast Vma12p and Vma22p. **B,** Mutations of VMA21 associated with X-linked myopathy with excessive autophagy and follicular lymphoma that map onto yeast Vma21p. Yeast residues are indicated with disease causing human mutations in brackets.

**Figure S8.**
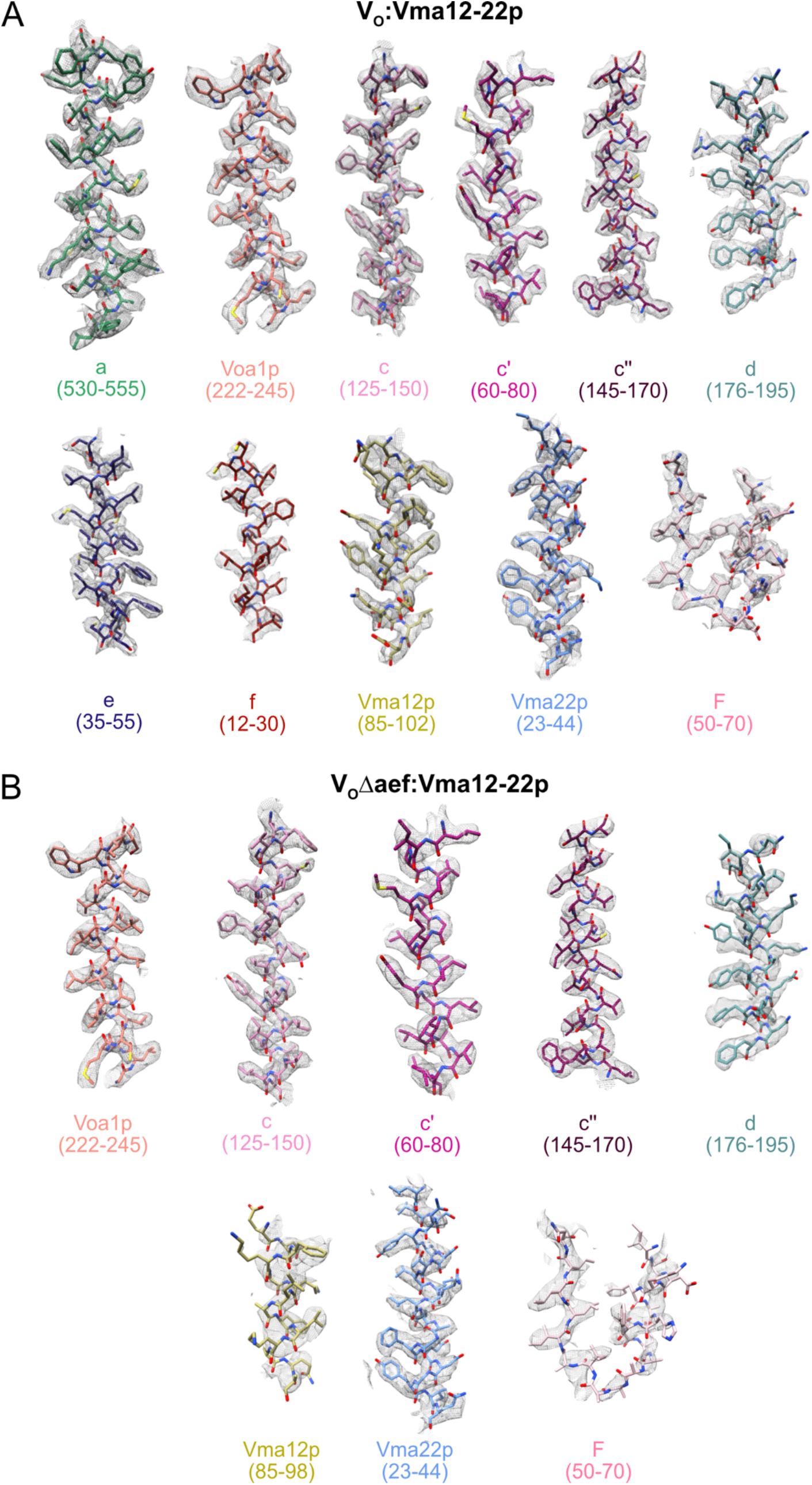
Model-in-map fit examples. Examples of the atomic model fit in the cryoEM map of V_O_:Vma12-22p (**A**) and V_O_Δaef:Vma12-22p (**B**).

**Figure S9.**
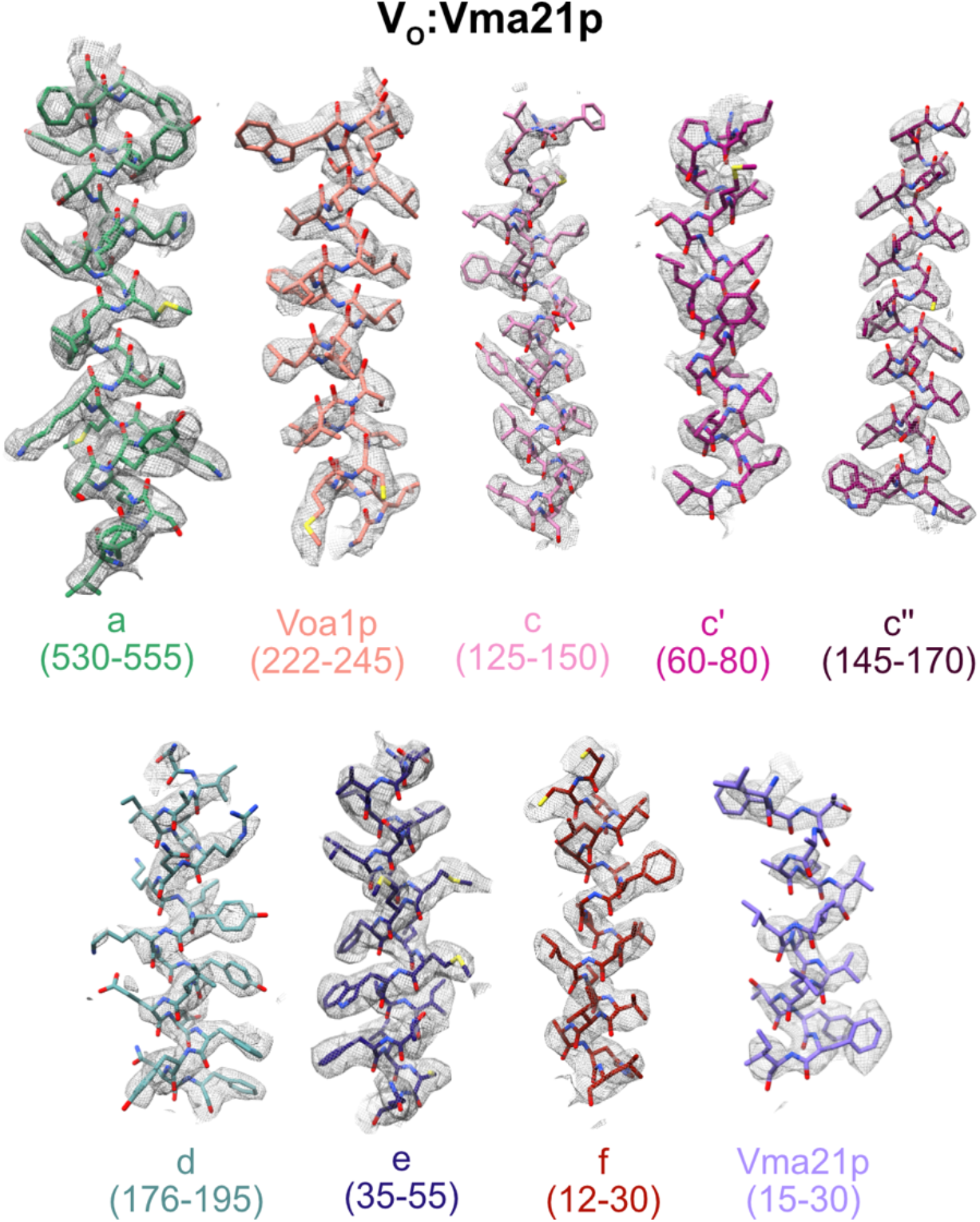
Model-in-map fit examples. Examples of the atomic model fit in the cryoEM map of V_O_:Vma21p.

**Table S1.**
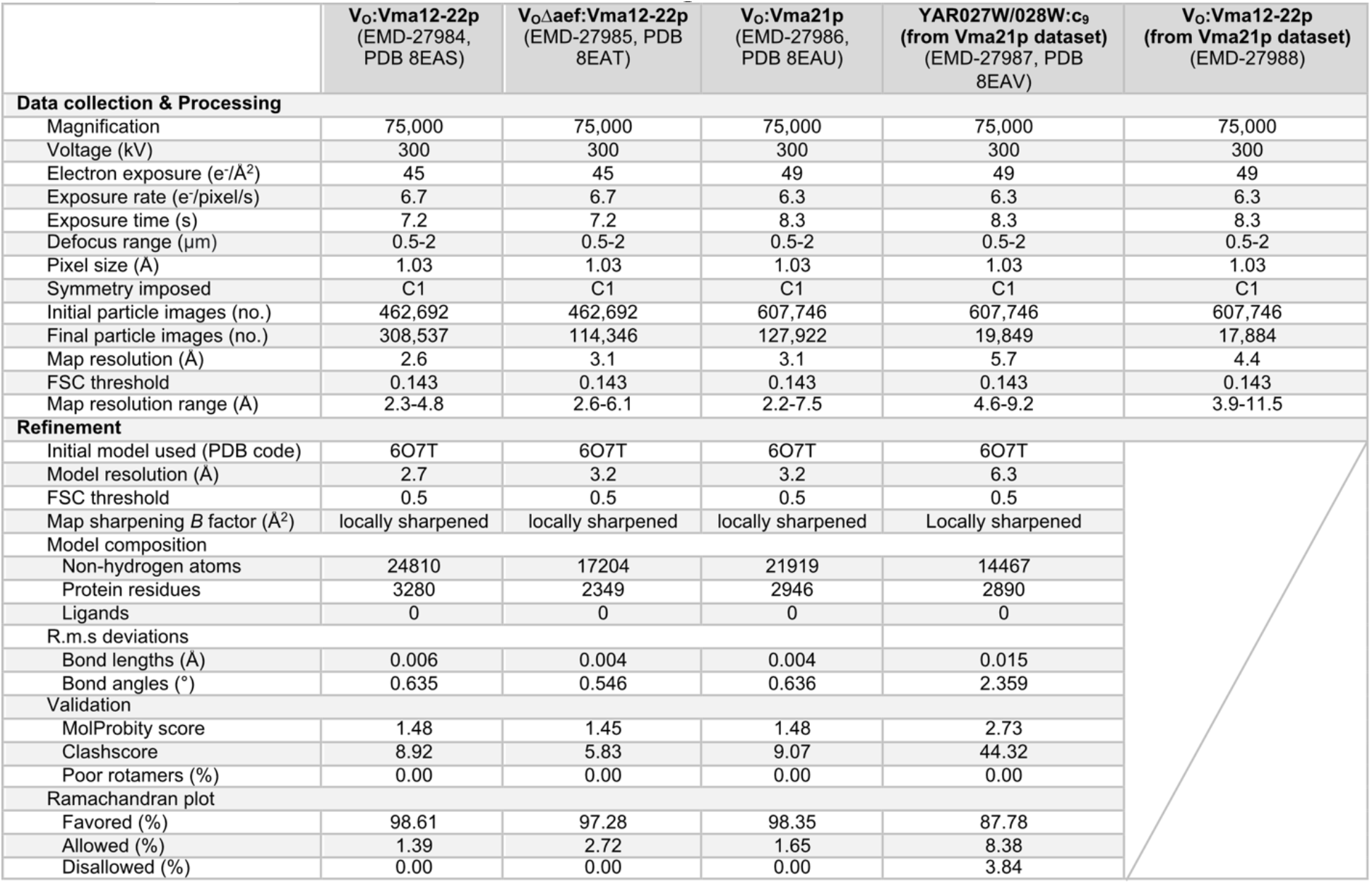
CryoEM and atomic model building statistics.

|  | V<sub>O</sub>:Vma12-22p<br>(EMD-27984,<br>PDB 8EAS) | V<sub>O</sub>Δaef:Vma12-22p<br>(EMD-27985, PDB<br>8EAT) | V<sub>O</sub>:Vma21p<br>(EMD-27986,<br>PDB 8EAU) | YAR027W/028W:c<sub>9</sub><br>(from Vma21p dataset)<br>(EMD-27987, PDB<br>8EAV) | V<sub>O</sub>:Vma12-22p<br>(from Vma21p dataset)<br>(EMD-27988) |
| --- | --- | --- | --- | --- | --- |
| Data collection & Processing |  |  |  |  |  |
| Magnification | 75,000 | 75,000 | 75,000 | 75,000 | 75,000 |
| Voltage (kV) | 300 | 300 | 300 | 300 | 300 |
| Electron exposure (e/Å<sup>2</sup>) | 45 | 45 | 49 | 49 | 49 |
| Exposure rate (e/pixel/s) | 6.7 | 6.7 | 6.3 | 6.3 | 6.3 |
| Exposure time (s) | 7.2 | 7.2 | 8.3 | 8.3 | 8.3 |
| Defocus range (μm) | 0.5-2 | 0.5-2 | 0.5-2 | 0.5-2 | 0.5-2 |
| Pixel size (Å) | 1.03 | 1.03 | 1.03 | 1.03 | 1.03 |
| Symmetry imposed | C1 | C1 | C1 | C1 | C1 |
| Initial particle images (no.) | 462,692 | 462,692 | 607,746 | 607,746 | 607,746 |
| Final particle images (no.) | 308,537 | 114,346 | 127,922 | 19,849 | 17,884 |
| Map resolution (Å) | 2.6 | 3.1 | 3.1 | 5.7 | 4.4 |
| FSC threshold | 0.143 | 0.143 | 0.143 | 0.143 | 0.143 |
| Map resolution range (Å) | 2.3-4.8 | 2.6-6.1 | 2.2-7.5 | 4.6-9.2 | 3.9-11.5 |
| Refinement |  |  |  |  |  |
| Initial model used (PDB code) | 6O7T | 6O7T | 6O7T | 6O7T |  |
| Model resolution (Å) | 2.7 | 3.2 | 3.2 | 6.3 |  |
| FSC threshold | 0.5 | 0.5 | 0.5 | 0.5 |  |
| Map sharpening *B* factor (Å<sup>2</sup>) | locally sharpened | locally sharpened | locally sharpened | Locally sharpened |  |
| Model composition |  |  |  |  |  |
| Non-hydrogen atoms | 24810 | 17204 | 21919 | 14467 |  |
| Protein residues | 3280 | 2349 | 2946 | 2890 |  |
| Ligands | 0 | 0 | 0 | 0 |  |
| R.m.s deviations |  |  |  |  |  |
| Bond lengths (Å) | 0.006 | 0.004 | 0.004 | 0.015 |  |
| Bond angles (°) | 0.635 | 0.546 | 0.636 | 2.359 |  |
| Validation |  |  |  |  |  |
| MolProbity score | 1.48 | 1.45 | 1.48 | 2.73 |  |
| Clashscore | 8.92 | 5.83 | 9.07 | 44.32 |  |
| Poor rotamers (%) | 0.00 | 0.00 | 0.00 | 0.00 |  |
| Ramachandran plot |  |  |  |  |  |
| Favored (%) | 98.61 | 97.28 | 98.35 | 87.78 |  |
| Allowed (%) | 1.39 | 2.72 | 1.65 | 8.38 |  |
| Disallowed (%) | 0.00 | 0.00 | 0.00 | 3.84 |  |

**Table S2. Mass spectrometry identification of proteins purified with Vma21p-3×FLAG preparation.**

**Supplementary Video S1. Model for the sequence of V_O_ assembly event.**

